# Efficient isolation of rare B cells using next-generation antigen barcoding

**DOI:** 10.1101/2022.06.06.495029

**Authors:** Jonathan Hurtado, Claudia Flynn, Jeong Hyun Lee, Eugenia Salcedo, Christopher A. Cottrell, Patrick D. Skog, David Nemazee, William R. Schief, Elise Landais, Devin Sok, Bryan Briney

**Affiliations:** Department of Immunology and Microbiology, Scripps Research, La Jolla, CA 92037, USA; Center for Viral Systems Biology, Scripps Research, La Jolla, CA 92037, USA; Center for HIV/AIDS Vaccine Development, Scripps Research, La Jolla, CA 92037, USA; International AIDS Vaccine Initiative (IAVI), New York, NY 10004, USA; IAVI Neutralizing Antibody Center, Scripps Research, La Jolla, CA 92037, USA; Ragon Institute of MGH, MIT and Harvard, Cambridge, MA 02139, USA; San Diego Center for AIDS Research, Scripps Research, La Jolla, CA 92037, USA

## Abstract

The ability to efficiently isolate antigen-specific B cells in high throughput will greatly accelerate the discovery of therapeutic monoclonal antibodies (**mAbs**) and catalyze rational vaccine development. Traditional mAb discovery is a costly and labor-intensive process, although recent advances in single-cell genomics using emulsion microfluidics allow simultaneous processing of thousands of individual cells. Here we present a streamlined method for isolation and analysis of large numbers of antigen-specific B cells, including next generation antigen barcoding and an integrated computational framework for B cell multi-omics. We demonstrate the power of this approach by recovering thousands of antigen-specific mAbs, including the efficient isolation of extremely rare precursors of VRC01-class and IOMA-class broadly neutralizing HIV mAbs.

## INTRODUCTION

The antibody repertoire is exceptionally diverse, allowing the humoral immune system to recognize and respond to a broad range of invading pathogens. The pre-immune antibody repertoire is generated by somatic recombination of variable (**V**), diversity (**D**) and joining (**J**) immunoglobulin gene segments, which occurs independently in each developing B cell [1]. In humans, it is estimated that the recombination process is capable of generating as many as 10^18^ unique antibody molecules [2]. Following antigen recognition, antibodies are affinity matured through an iterative process of clonal expansion, somatic hypermutation, and antigen-driven selection [3,4]. Following pathogen clearance, a subset of B cells encoding affinity matured antibodies are retained as an immune “memory” of the pathogen encounter. Humoral immune memory, which can persist for decades [5], rapidly reactivates in response to subsequent exposure to the same pathogen and is the primary mechanism of protection for most available vaccines [6,7]. Monoclonal antibodies (**mAbs**) are invaluable tools for the treatment and prevention of human disease. Antigen-specific mAbs are useful as templates for rational vaccine development, in which immunization strategies are designed to preferentially elicit antibodies encoding a defined set of genetic or structural properties [8–11]. Additionally, passively delivered therapeutic mAbs have a variety of clinical applications, including cancer, autoimmunity and infectious disease [12,13].

Exceptionally potent neutralizing antibodies (**nAbs**) are often quite rare and may be found only in a subset of seropositive individuals, meaning their discovery typically requires deep interrogation of the pathogen-specific B cell repertoire [14]. Traditional techniques for isolating antigen-specific human mAbs are expensive and immensely labor intensive; despite these obstacles, mAb-based therapies against emerging infectious diseases have a distinct advantage in that their discovery and clinical advancement may proceed more quickly than traditional small molecule drugs. This was highlighted during the COVID-19 pandemic, in which clinical trials of novel mAb-based therapeutics were initiated just months after the first known cases of SARS-CoV-2 infection [15–17]. Additionally, broadly neutralizing antibodies (**bnAbs**) with the ability to recognize a wide range of viral variants or even entire families of related viruses [18–29] are also useful as templates to guide rational vaccine development strategies by revealing conserved sites of viral vulnerability [30,31].

Recovery of natively paired mAb sequences is most commonly performed by sequestering individual B cells prior to amplifying and sequencing the heavy and light chains from each cell. There are various methods of sequestration, including limiting dilution of immortalized or transiently activated primary B cells [23,32–34], deposition of single B cells into discrete wells [35–37], or in-cell amplification techniques in which the cell itself serves as the encapsulation vessel [38,39]. These processes are immensely costly and labor intensive, meaning even large-scale studies are often only able to isolate dozens or hundreds of mAbs with the desired specificity. Recent advances in emulsion microfluidics have dramatically increased the scale at which cellular sequestration can be performed, removing a significant bottleneck in the mAb discovery process and enabling routine recovery of up to thousands of natively paired mAbs in a single experiment. Indeed, the largest single collection of natively paired mAb sequences, containing sequences from 1.6×10^6^ single B cells, was recently obtained using the emulsion microfluidics-based 10x Genomics Chromium X platform [40].

In 2017, CITE-seq (***C***ellular ***I***ndexing of ***T***ranscriptomes and ***E***pitopes by ***seq***uencing) pioneered the use of DNA-barcoded antibodies to simultaneously quantify transcription and protein expression using single cell droplet microfluidics [41]. Briefly, antibodies against a defined panel of cell surface markers were tagged with a unique DNA barcode and used to label cells. The barcoding oligonucleotides contain an antibody-specific barcode, a unique molecular identifier (**UMI**) and sequencing platform-specific primer annealing sites. During single cell library preparation, barcodes and cellular mRNAs are recovered and sequenced, and the UMI-normalized sequencing read counts can be used to quantify both transcription and protein abundance. The available number of distinct CITE-seq barcodes is effectively unlimited, allowing the use of much larger antibody panels than would be feasible with even the most sophisticated flow cytometers currently available [42]. Building on the technical and conceptual foundation established by CITE-seq, approaches for linking adaptive immune receptor sequences with antigen specificity were developed, first for T cell receptors (**TCRs**) and subsequently for B cell receptors (**BCRs**) [43,44]. LIBRA-seq (***LI***nking ***B*** cell ***R***eceptor and ***A***ntigen through ***seq***uencing) enables simultaneous assessment of BCR sequence and specificity using DNA-barcoded antigens [44]. By modifying the CITE-seq and TCR antigen specificity approaches that preceded it, LIBRA-seq entails labeling antigen-specific B cells with one or more DNA-barcoded antigens prior to single cell library generation using the 10x Genomics platform, thus linking BCR sequence and specificity (Figure 1a). Like CITE-seq, the available number of unique antigen barcode sequences is not a limiting factor, however, the number of antigen baits which compete for BCR binding will be somewhat constrained by the finite number of BCR molecules on the cell surface and the need to ensure the signal of each antigen bait is distinguishable from background.

**Figure 1.**
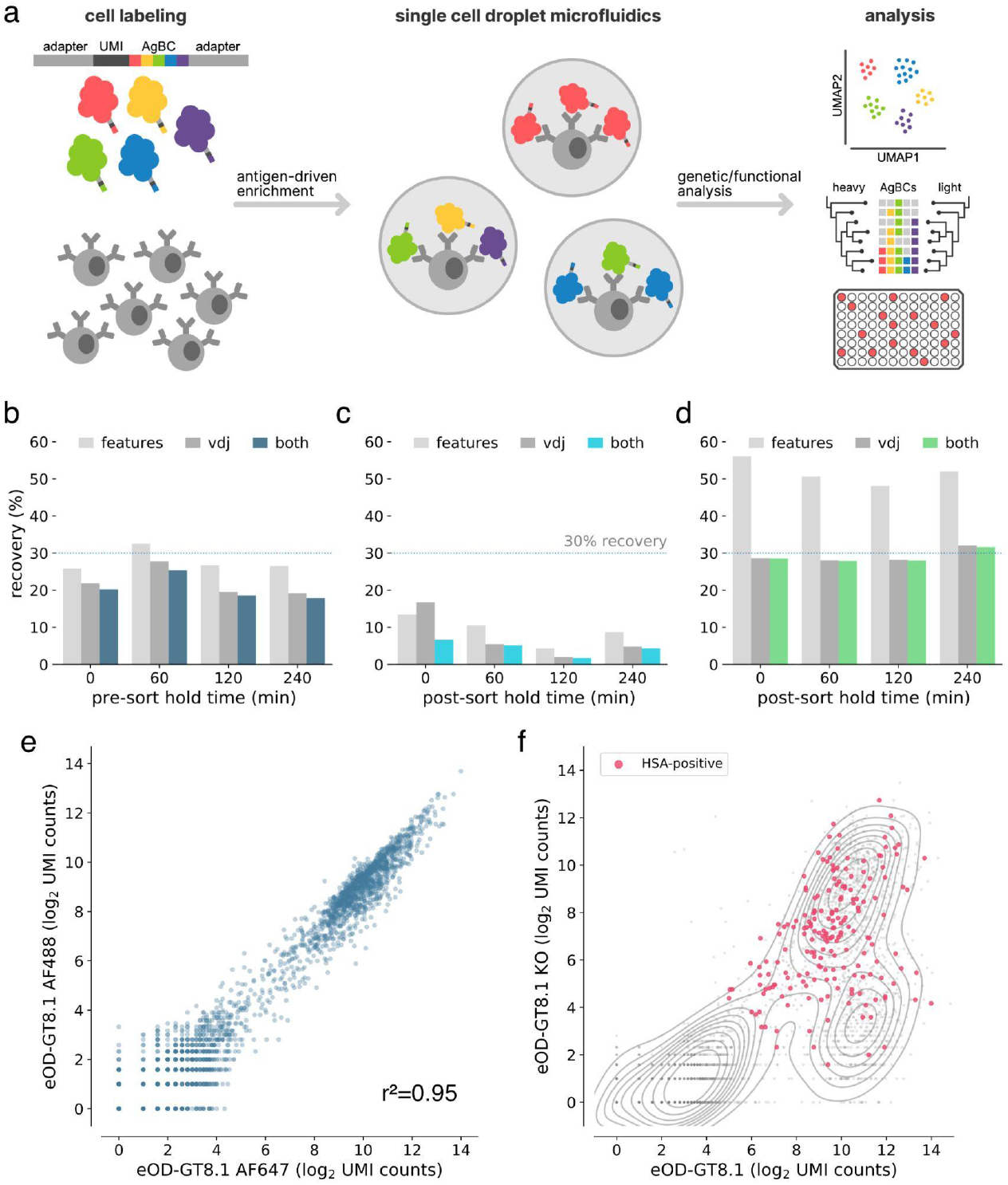
Optimization of next-generation AgBCs. (**a**) Schematic overview of the antigen barcoding approach. (**b**) Pre-optimization recovery of features (light gray), paired VDJ sequences (dark gray) or both features and VDJ (blue) when sorting cells after pre-sort hold times of varying duration. (**c**) Pre-optimization recovery of features (light gray), paired VDJ sequences (dark gray) or both features and VDJ (blue) when processing sorted cells after post-sort hold times of varying duration. (**d**) Optimized recovery of features (light gray), paired VDJ sequences (dark gray) or both features and VDJ (green) when processing sorted cells after post-sort hold times of varying duration. (**e**) Correlation between recovery AgBC UMI counts of two differently barcoded aliquots of eOD-GT8.1. Plot was generated in scab using scab.pl.feature_scatter(). (**f**) Kernel density estimate plot of the log2-normalized UMI counts of eOD-GT8.1 AgBC and eOD-GT8.1 KO AgBC is shown in gray. Cells classified as positive for an HSA AgBC are highlighted in pink. Plot was generated in scab using scab.pl.feature_kde().

Here we present an optimized version of the original LIBRA-seq technique, including the development of next-generation antigen barcodes (**AgBCs**), as a streamlined method for isolating antigen-specific B cells. We also report the development of a computational framework for large-scale analysis of B cell-derived single cell omics data, which provides tools for multi-omics data integration, accurate assignment of BCR specificity using AgBCs, clonal lineage assignment, and visualization. Finally, we demonstrate the power of this improved approach by simultaneously isolating multiple classes of extremely rare HIV broadly neutralizing antibody (**bnAb**) precursors from HIV-uninfected donors using a panel of germline targeting immunogens.

## RESULTS

### Optimizing experimental conditions to maximize recovery of FACS-enriched live B cells

Discovering rare antibodies often requires antigen-driven enrichment using samples containing many millions of B cells, which translates into prolonged fluorescence activated cell sorting (**FACS**) experiments lasting several hours. A critical factor affecting sequence recovery from the 10x Genomics platform is cell viability. In the original LIBRA-seq study, paired BCR sequences and antigen barcodes were recovered from 6-22% of input cells (see *Methods and Methods* for details about the LIBRA-seq recovery efficiency calculations). We hypothesized that the incubations required during extended FACS experiments, either pre-sort (queued cells waiting to be sorted) or post-sort (cells already sorted but awaiting completion of the experiment) contributed to the low yield and high variability in recovery between samples. To test this, splenocytes from four C57BL/6 mice and four VRC01 gH mice, which express a germline-reverted heavy chain from the HIV bnAb VRC01 [8,45], were individually labeled with unique cell hashes and pooled. Aliquots of the hashed splenocyte pool were either (***1***) held on ice for varying intervals (0, 1, 2 or 4 hours) prior to sorting and library generation using a 10x Genomics Chromium Controller, or (***2***) immediately sorted and held on ice for varying intervals post-sort (0, 1, 2, or 4 hours) before proceeding to library generation. For each aliquot, 1×10^5^ random CD19+ cells were sorted and used to prepare BCR and feature barcode sequencing libraries. We observed no difference in recovery for samples subjected to a pre-sort hold, but noted a substantial decline in recovery efficiency for samples held on ice after sorting (Figure 1b-c). Notably, recovery using the unoptimized protocol closely matched the range of efficiencies in the original LIBRA-seq protocol. We evaluated a variety of experimental conditions to identify optimal parameters that maximize recovery of B cells held for several hours following FACS enrichment. These included cell fixation using paraformaldehyde or methanol, sorting into tubes and plates of varying sizes and types, and systematically assessing catch buffer volume and composition (data not shown). In the end, we found that the optimal conditions involved bulk sorting unfixed cells into a 96-well PCR plate well containing a low volume catch buffer of 20μL of 100% fetal bovine serum (**FBS**). Using these conditions, we consistently obtained high recovery efficiency (>25%) regardless of the post-sort hold duration (Figure 1d).

### Streamlined production of next-generation DNA barcoded antigens

We sought to improve the LIBRA-seq method by enhancing the antigen barcode complexes (**AgBCs**) used to link BCR sequence and specificity. Previously, AgBCs were generated in a multi-step process consisting of (***1***) site-specific biotinylation of protein antigens, (***2***) direct conjugation of barcode oligonucleotides to the biotinylated antigen, and (***3***) addition of a streptavidin-linked fluorophore to the barcoded and biotinylated protein antigen. This approach has a significant drawback in that the oligonucleotide conjugation is not performed in a site-specific manner. As a result, critical B cell epitopes are likely to be occluded or disrupted by the barcode oligonucleotide. The original LIBRA-seq publication also cautions that the direct oligonucleotide conjugation approach may result in substantial heterogeneity in the number of barcode oligonucleotides attached to each antigen molecule, particularly across different antigens and between different conjugation batches of the same antigen, which may confound downstream analyses [44]. We have simplified the construction of next-generation AgBCs through the use of a barcoding reagent generated by incubating a fluorophore-linked streptavidin (or plain streptavidin, if a fluorophore is not desired) with a 5’-biotinylated barcode oligonucleotide. Importantly, the biotinylated oligonucleotide is added at a sub-saturating concentration (2.5:1 molar ratio of oligonucleotide:streptavidin), which ensures that the majority of the resulting barcoding reagent molecules will have unoccupied biotin binding sites as well as a similar number of barcode oligonucleotides. The barcoding reagent is then incubated with a site-specifically biotinylated protein antigen. Each antigen molecule is thus linked to the barcoding reagent in a site-specific manner that minimizes alterations to the native structure of the antigen.

### Selective enrichment of antigen-specific B cells using AgBCs

Dual antigen labeling strategies, in which aliquots of the same antigen are conjugated to different fluorophores and used to select B cells positive for both fluorescent markers, are commonly used to reduce background when performing antigen-driven enrichment of B cells [46,47]. By constructing two AgBCs for each antigen, each with a different fluorophore and barcode, we can reduce background and improve enrichment selectivity both during sorting using dual fluorophores and post-sort data analysis using dual barcodes. To evaluate the reproducibility of our AgBCs, we stained B cells from transgenic mice expressing the human IGHV1-2*02 variable gene (huVH1-2 KI mice, [48]) with differently barcoded aliquots of the engineered HIV immunogen eOD-GT8.1 [49], which is designed to target germline precursors of VRC01-class HIV bnAbs. UMI-normalized read counts of the differently barcoded eOD-GT8.1 AgBCs were well correlated (r^2^=0.95), indicating a high level of reproducibility (Figure 1e). In each of our mAb discovery experiments, we also include a dark (lacking a fluorophore) human serum albumin (**HSA**) AgBC. This negative control AgBC, which is not detectable during sorting, allows us to identify and remove cells during downstream data analysis to which AgBCs are binding in a nonspecific manner. This is particularly important when assessing BCR breadth against a panel of AgBCs, as nonspecific binding to “sticky” cells can mimic the expected binding patterns of broadly reactive BCRs. Demonstrating the importance of including a negative control AgBC, a representative sample of B cells from VRC01 gH mice sorted using two competing AgBCs (eOD-GT8.1 and eOD-GT8.1 KO, an epitope knockout variant that disrupts binding of antibodies that recognize the CD4 binding site), a substantial fraction of cells that appeared to specifically bind both eOD-GT8.1 and eOD-GT8.1 KO were also positive for HSA (Figure 1f). HSA+ and HSA-B cells were not distinguishable based solely on binding to eOD-GT8 and eOD-GT8 KO, emphasizing the importance of control AgBCs to assigning specificity profiles with high confidence.

### Computational analysis and integration of single B cell multi-omics data

To process and integrate the various datasets obtained by single cell sequencing using AgBCs and the 10x Genomics platform, which can include gene expression (**GEX**), paired BCR and TCR sequences, AgBCs, cell hashes and other feature barcodes such as CITE-seq antibodies, we have developed *scab*, a Python package for ***S***ingle ***C***ell ***A***nalysis of ***B*** cells. We developed scab primarily as a means to perform interactive analyses in combination with notebook-like programming environments such as Jupyter [50,51]. To integrate these datasets, scab utilizes the Sequence and Pair objects introduced in our ab[x] toolkit for antibody sequence analysis [52] to link annotated BCR/TCR sequence information with GEX, antigen specificity, and feature barcode (**FBC**) data for each cell in an integrated AnnData object [53]. Scab includes utilities to read and integrate data produced by CellRanger [54] and write integrated AnnData objects as .h5ad-formatted hdf5 files. During data ingestion, scab will automatically annotate BCR and TCR contigs using abstar [52]. Scab also includes a variety of tools for interactive single cell data analysis and exploration, including sample demultiplexing using cell hashes, BCR specificity classification using AgBCs, and clonal lineage assignment using the clonify algorithm [55], as well as sophisticated visualization tools for exploratory data analyses and generating publication-quality figures. An illustrated code example is shown in Figure 2, which performs a common sequence of operations: data ingestion and automatic BCR sequence annotation, demultiplexing, specificity classification, clonal lineage assignment and saving the fully annotated AnnData object to disk as an h5ad-formatted file.

**Figure 2.**
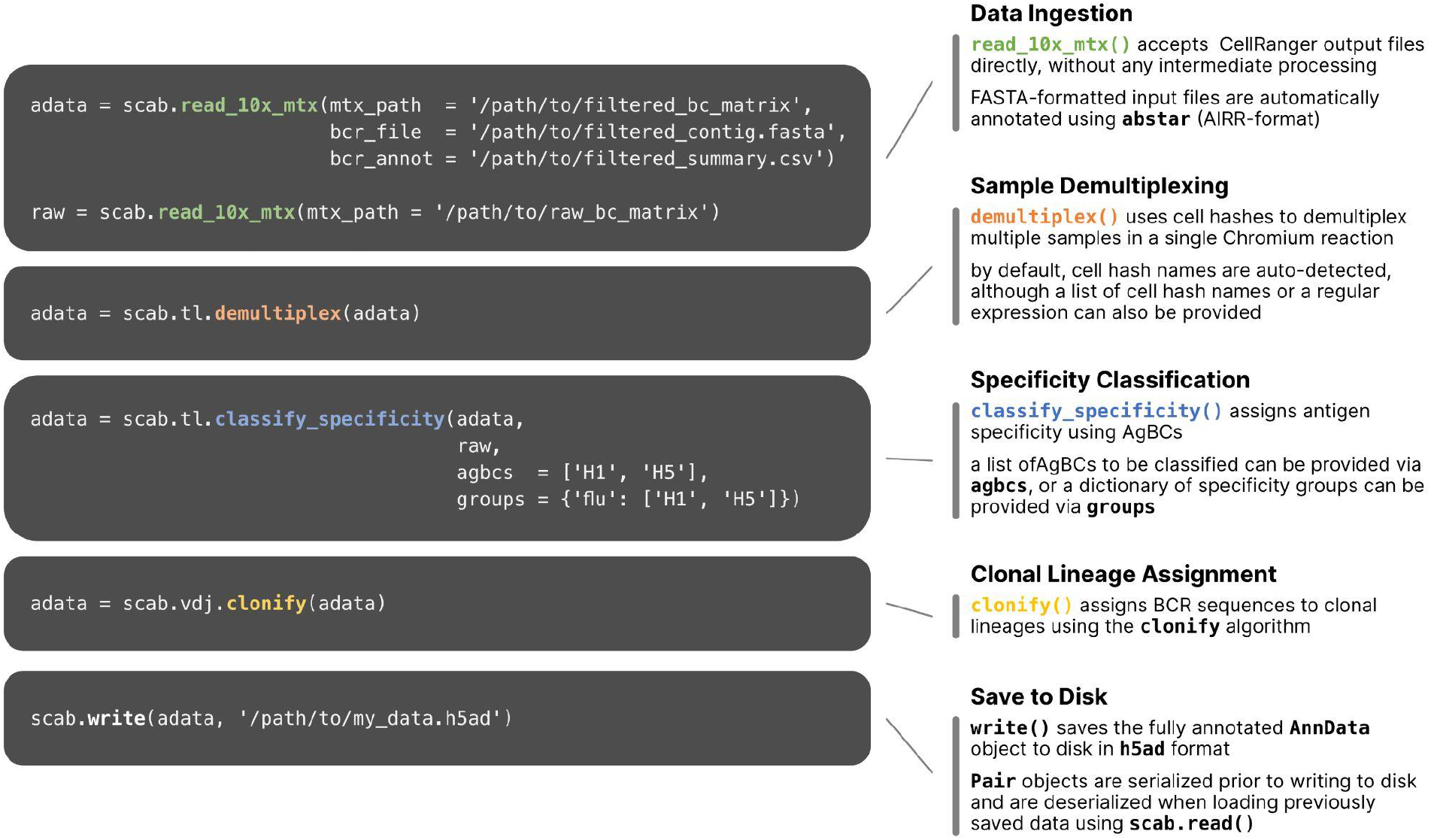
Workflow of single cell multi-omics data processing using scab. Example commands are shown for several common data processing and analysis tasks. Briefly, starting with the unmodified output files from CellRanger, this sample workflow (**1**) reads CellRanger output data and annotates BCR sequences using abstar [52]; (**2**) demultiplexes samples using cell hashes; (**3**) classifies BCR specificity using two AgBCs (“H1” and “H2”) and a single specificity group (“flu”), which includes both AgBCs; (**4**) assigns BCR sequences to clonal lineages using clonify [55]; (**5**) writes the integrated AnnData object to file in h5ad format.

Tools for cell hash-based sample demultiplexing have previously been reported [56,57], however, our experience with existing methods based on Gaussian mixture models (**GMMs**) or k-means clustering have shown reduced accuracy when the distribution of hashes is highly unequal (that is, when some hashes are found much more or much less frequently than others). Specifically, we noted degraded performance of GMM-based cell hash classification for individual samples comprising less than 5% of the overall sample pool. Because the frequency of antigen-specific B cells can vary substantially between individuals, we often multiplex samples that differ greatly in cell number. Thus, we sought to develop an alternative approach designed to accurately demultiplex highly heterogeneous cell hash pools. For each cell hash, we compute a kernel density estimate (**KDE**) of the log_2_-transformed, UMI-normalized read counts. We then identify the local minimum that optimally separates the KDE into two populations: positive and negative (Figure 3a). This inflection point is used as the cell hash classification threshold. This approach accurately demultiplexes samples even when the frequency of individual cell hashes differs by nearly an order of magnitude (Figure 3b). Cells that exceed the threshold for multiple hashes are considered doublets, and cells that do not exceed any threshold are considered unassigned (Figure 3c).

**Figure 3.**
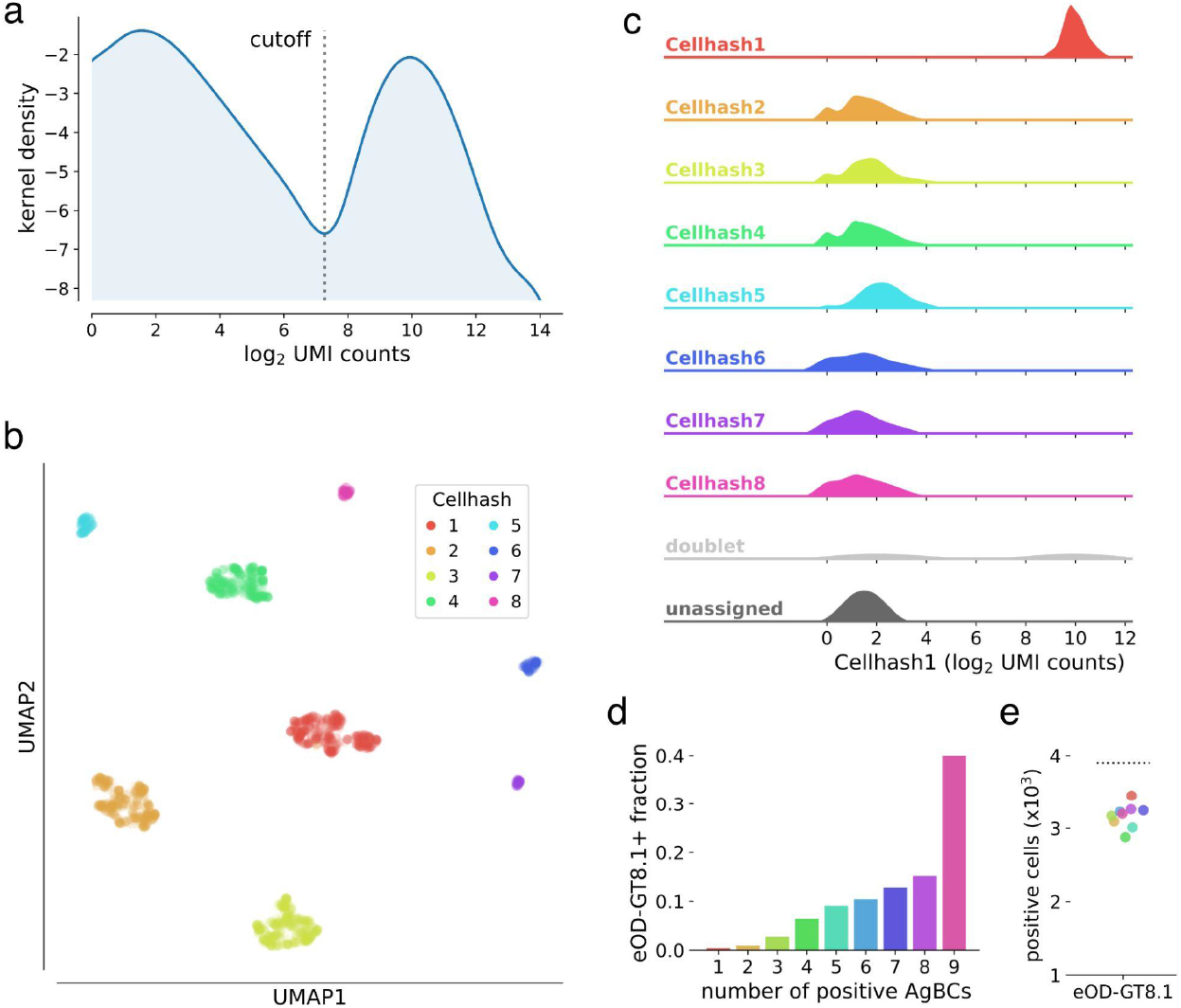
Classification of cell hashes and AgBCs. (**a**) Representative KDE plot of log_2_-transformed cell hash UMI counts. The classification cutoff is highlighted, and clearly separates the positive and negative cell hash populations. KDE plots are optionally generated when running scab.tl.demultiplex(). (**b**) UMAP plot of cells clustered using log_2_-transformed cell hash UMI counts. Cells are colored by cell hash assignment. Of note, cell hashes 1-4 each comprise 20-25% of all input cells, while cell hashes 5-8 each comprise 2-5% of all input cells. (**c**) Ridgeplot showing the distribution of log_2_-transformed UMI counts of cell hash 1 for all cell hash assignment groups. Plot was generated with scab by calling scab.pl.cellhash_ridge(). (**d**) Fraction of cells classified as positive for increasing numbers of differently barcoded aliquots of eOD-GT8.1 AgBC when performing classification on each AgBC separately. A total of 9 different AgBCs were used, meaning approximately 40% of all recovered cells were classified as positive for all AgBCs. (**e**) Fraction of cells classified as positive for each of 9 differently barcoded eOD-GT8.1 AgBCs when classification was performed separately for each AgBC (colored points). The dashed line shows the number of cells classified as eOD-GT8.1 positive when using a specificity group containing all 9 eOD-GT8.1 AgBCs.

Classification of BCR specificity using AgBCs is made more complex by the potentially wide range of binding affinities displayed by antigen-specific BCRs and a variable number of membrane-bound BCR molecules on the cell surface of each B cell. These factors combine to produce a gradient of UMI-normalized count values for each AgBC rather than the clearly defined positive and negative populations typical of CITE-seq or cell hash data (Figure 3a). Separately for each AgBC, we first inspect “empty” droplets in which CellRanger did not identify a cell. We then compute a negative binomial distribution of UMI-normalized AgBC counts and establish the background threshold at the 99.7th percentile (3σ). Any cell-containing droplets with UMI-normalized AgBC counts above this threshold are considered antigen positive. When working with large AgBC panels containing antigens that may compete for BCR binding (for example, epitope mapping panels or panels containing multiple antigen variants to identify cross-reactive BCRs), it is possible that competition for the finite number of BCR molecules on the B cell surface may reduce the number of bound AgBCs for one or more of the competing antigens, resulting in incorrect specificity classification. To combat this, we introduced the concept of specificity groups, which represent a group of AgBCs that are expected to compete for BCR binding. When performing specificity classification, the classification threshold is set by considering the sum of all grouped AgBC UMI counts in empty droplets, and any cell-containing droplets whose summed AgBC UMI counts exceeding the threshold are considered positive. To demonstrate the utility of this approach, we separately barcoded nine aliquots of eOD-GT8.1 and sorted antigen-positive B cells using the pool of eOD-GT8.1 AgBCs. While approximately 40% of all sorted cells were classified as positive for all nine AgBCs individually (Figure 3d), the use of a specificity group containing all nine AgBCs correctly classified a significantly higher number of cells than did any of the competing AgBC individually (Figure 3e).

### Isolation of ultra-rare HIV broadly neutralizing antibody precursors

The design of vaccine immunogens that reliably activate unmutated bnAb precursors and guide their development toward mature bnAbs is the foundation of germline-targeting and epitope-focusing vaccine development strategies [58]. For HIV in particular, bnAbs tend to require uncommon genetic features and their precursors are expected to be found at very low frequencies in the circulating naive B cell repertoire [59–61]. VRC01-class HIV bnAbs, which all encode a conserved set of required genetic features including the IGHV1-2 heavy chain variable gene and a short (5 amino acid [AA]) light chain complementarity determining region 3 (**LCDR3**), target the CD4 binding site (**CD4bs**) on HIV Env and are among the most broad and potent classes of HIV bnAbs [62]. Although inferred germline variants of VRC01-class bnAbs typically do not detectably bind HIV, germline targeting immunogens have been designed which selectively target naive B cell precursors which encode the genetic features required of VRC01-class bnAbs [8,10,49,63,64]. Interrogating the B cell repertoires of HIV-uninfected individuals with these immunogens revealed the frequency of VRC01-class bnAb precursors to be roughly 1 in 3×10^5^-2.4×10^6^ circulating naive B cells [49,65]. IOMA-class HIV bnAbs are similar to VRC01-class bnAbs in that they also target the CD4bs and use IGHV1-2, but IOMA-class light chains using IGLV2-23 and encode an 8 AA LCDR3 [66]. Although less broad and potent than VRC01-class bnAbs, it has been hypothesized that IOMA-class bnAbs will be easier to elicit by vaccination due to its lower level of somatic mutation and the use of a much more common LCDR3 length.

We constructed two separately barcoded AgBCs for eOD-GT8.1 and eOD-GT8.1 KO. Combined with a dark HSA AgBC and a dually barcoded positive control antigen (BG505 SOSIP), the complete AgBC panel was used to isolate antigen-specific B cells from a total of 7.04×10^8^ peripheral blood B cells from two healthy adult donors (D931 and D326651). We processed a total of 24,792 sorted cells across two Chromium 5’ Single Cell v2 reactions and recovered 9,216 paired antibody sequences for an overall paired sequence recovery efficiency of 37.2%. After AgBC processing and removal of “sticky” cells that displayed substantial HSA AgBC binding, we recovered a total of 2,618 eOD-GT8.1-specific B cells. Of these, 162 B cells were classified as positive for eOD-GT8.1 and negative for eOD-GT8.1-KO, indicating epitope specific binding to the HIV CD4bs. From the eOD-GT8.1 positive population, we identified 18 VRC01-class B cells (encoding IGHV1-2 and a 5AA LCDR3) and four IOMA-class B cells (encoding IGHV1-2, IGLV2-23, and an 8AA LCDR3). Notably, all of the VRC01-class and IOMA-class B cells displayed AgBC binding patterns consistent with CD4bs specificity (Figure 4a). Our VRC01-class recovery equates to an overall frequency of 1 in ∼3.9×10^6^ B cells (unadjusted for expected losses during sorting and library preparation), which is consistent with other unadjusted precursor frequency estimates [49,65,67]. Our recovery of IOMA-like precursors mirrors previous work using eOD-GT8.1 tetramers and 60-mers [65].

**Figure 4.**
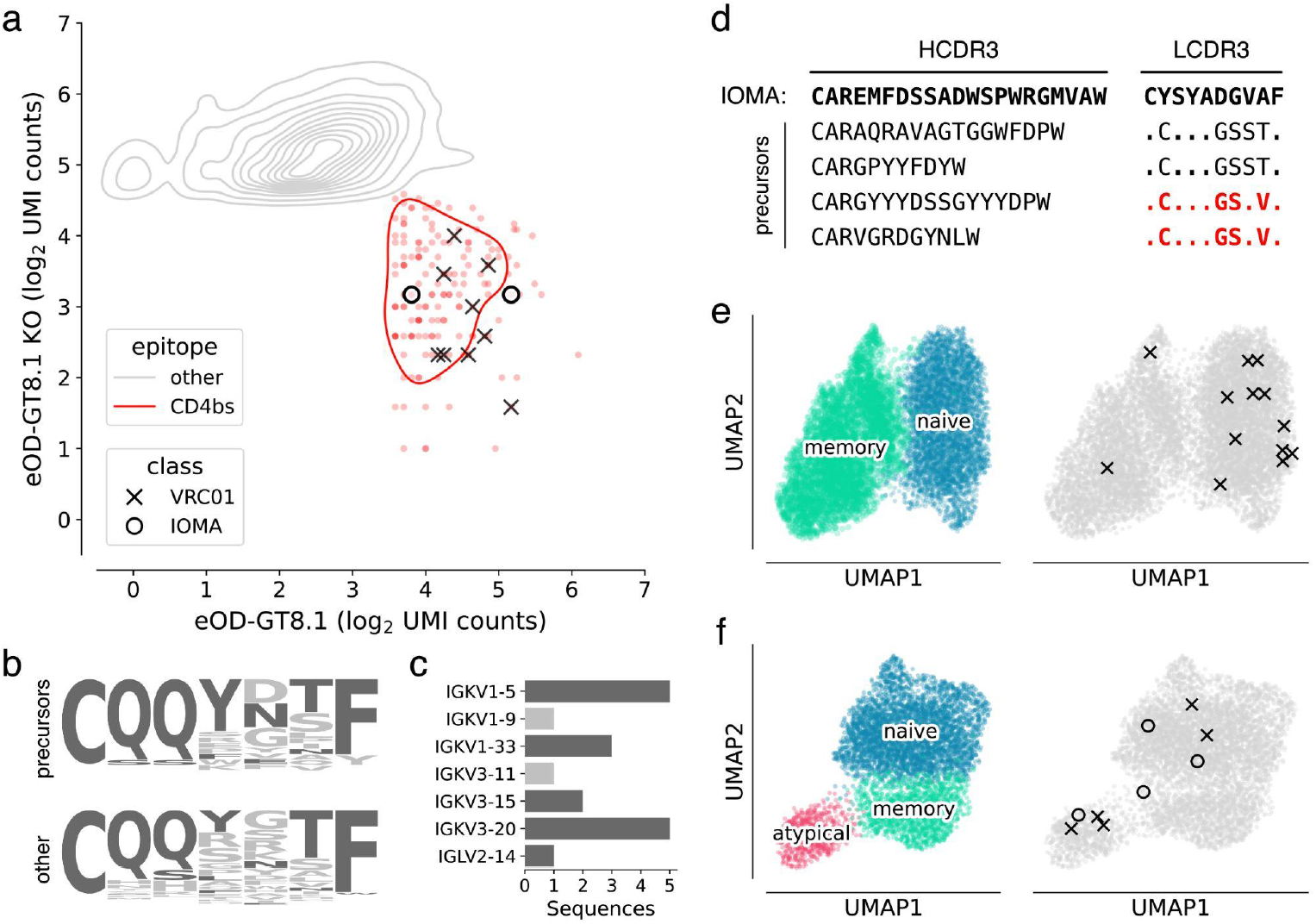
Characterization of VRC01-class and IOMA-class HIV bnAb precursors. (**a**) Density plot of log_2_-normalized AgBC UMI counts for eOD-GT8.1 (x-axis) and eOD-GT8.1 KO (y-axis). Cells positive for eOD-GT8.1 and negative for eOD-GT8.1 KO (indicating binding to the CD4bs epitope) are indicated in red, all others are in gray. VRC01-class (X) and IOMA-class (O) are highlighted. Plot was generated with scab by calling scab.pl.feature_kde(). (**b**) LCDR3 logo plots for all recovered VRC01-class precursors (top) and all other 5AA LCDR3s, which were not paired to a heavy chain encoding IGHV1-2 (bottom). Residues found in mature VRC01-class bnAbs are colored dark gray, all other residues are light gray. (**c**) Light chain V-gene use in isolated VRC01-class precursors. Light chain V-genes used by mature VRC01-class bnAbs are colored dark gray, all others are colored light gray. (**d**) Heavy chain and light chain CDR3 sequences for isolated IOMA-class precursors. LCDR3s that have been previously reported in IOMA-class precursors [65] are highlighted in red. (**e**) UMAP plots using single cell RNA-seq data for all recovered cells from D931. On the left, cells are colored by subset, with naive B cells in blue and memory B cells in green. The same UMAP embedding is plotted on the right in gray with cells encoding VRC01-class (X) or IOMA-class (O) precursors highlighted. (**f**) UMAP plots using single cell RNA-seq data for all recovered cells from D326651. On the left, cells are colored by subset, with naive B cells in blue, memory B cells in green, and atypical B cells in pink. The same UMAP embedding is plotted on the right in gray with cells encoding VRC01-class (X) or IOMA-class (O) precursors highlighted. Plots in (**e**) and (**f**) were generated with scab by calling scab.pl.umap().

The genetics of the isolated bnAb precursors (VRC01-class and IOMA-class) closely matched our expectations, based on the sequences of their respective bnAbs and prior studies which isolated CD4bs-targeting bnAb precursors with more traditional techniques. The LCDR3s of isolated VRC01-class precursors showed enrichment of residues found in mature VRC01-class bnAbs (Figure 4b) and 16 of 18 precursors encode light chain V genes used by mature VRC01-class bnAbs (Figure 4c). The two remaining light chain V genes, IGKV1-9 and IGKV3-11, have been observed in previously reported VRC01-class precursors isolated with eOD-GT8.1 [67]. As in previous studies, IOMA-class precursors encode diverse HCDR3s of varying lengths, but the critical LCDR3 region is well conserved (Figure 4d). Indeed, two of the LCDR3 sequences we recovered were identical to previously isolated IOMA-class bnAb precursors [65].

A significant advantage of our next-generation antigen barcoding approach compared to traditional techniques based on single cell sorting and PCR is the ability to simultaneously recover single cell transcriptional profiles together with full-length BCR sequences and AgBC data. This enables a more holistic analysis of the B cell repertoire, linking phenotypic properties like cellular activation and differentiation states with BCR genetics and antigen specificity. All of the VRC01-class and IOMA-class precursor sequences we isolated were completely unmutated in both heavy and light chains and were of the IgM isotype, suggesting they were recovered from naive B cells. Analysis of single cell transcriptomics data, however, shows that two of the B cells encoding VRC01-class precursors from donor D931 belong to the memory B cell subset (Figure 4e). Additionally, three VRC01-class precursors and one IOMA-class precursor from donor D326651 are encoded by atypical B cells (Figure 4f), a B cell subset that was recently shown to be more frequent in healthy individuals than previously thought [68].

## DISCUSSION

Efficient isolation of large numbers of pathogen-specific mAbs is critically important for fundamental immunological analyses of host/pathogen interactions, discovery of biological therapeutics for a variety of human diseases, and the development of efficacious vaccines. Here, we present an optimized protocol for mAb isolation using next-generation antigen barcoding as well as an integrated computational framework for analysis and visualization of B cell multi-omics data. We demonstrated the utility of our improved protocol by isolating thousands of natively paired mAb sequences specific for eOD-GT8.1, an HIV germline targeting immunogen, including several extremely rare VRC01-class and IOMA-class bnAb precursors. This streamlined approach removes a significant obstacle to the large-scale study of humoral immune responses to vaccination and infection.

As is common, however, removal of one experimental hurdle reveals the existence of yet another bottleneck further downstream in the process. In this case, streamlining the isolation of natively paired sequences for large numbers of antigen-specific mAbs results in sequence datasets that far exceed our capacity for biophysical characterization, including structural analyses, fine epitope mapping, and evaluation of additional functional properties beyond binding specificity (neutralization, effector function, protection, etc). In the future, it is possible that paradigm-shifting advances such as those seen recently with AlphaFold [69] will produce computational models capable of accurately inferring many or all of these functional properties directly from mAb sequences. Such models will be crucially important moving forward, even if they do not completely eliminate the need for recombinant expression and evaluation. One can easily envision a scenario in which large numbers of mAbs with desirable binding properties (isolated using techniques similar to those described here) can be screened *in silico* using models adept at predicting structure [70,71], fine epitope specificity [72–75], and other functional properties [76]. The results of *in silico* screening can be used to make informed down-selection decisions to focus limited biophysical characterization efforts toward the most interesting mAbs. As an example, structure and epitope predictions can be used to group similar antibodies and identify a relatively small number of representative mAbs that epitomize the functional properties of each group. Moving forward, we envision a virtuous cycle in which large mAb/specificity datasets are used to train better models and algorithms, which will in turn allow accurate analysis of larger or more complex mAb/specificity datasets. Increasingly rich datasets that accurately link natively paired antibody sequences with specificity information will be invaluable tools for training predictive models that may one day reduce or eliminate entirely the need for recombinant expression and characterization of large numbers of mAbs and unlock far more comprehensive studies of humoral immunology than are currently possible.

## MATERIALS AND METHODS

### Mouse samples

Three mouse models were used in this study: (***1***) VRC01-gH mice, which contain a transgene encoding the recombined, germline-reverted heavy chain of the human anti-HIV antibody VRC01 [8]; (***2***) huVH1-2 KI mice, which contain a transgene encoding the human variable gene IGHV1-2*02, which is a necessary genetic feature of the human anti-HIV antibody VRC01 [48]; and (***3***) C57BL/6J mice (Jackson Laboratories). Mouse studies were approved and carried out in accordance with the Institutional Animal Care and Use Committee at Scripps Research (La Jolla, CA). The mice were housed, immunized, and euthanized at Scripps in compliance with the Guide for the Care and Use of Laboratory Animals (National Research Council, 1996).

### Human samples

Human leukapheresis samples were obtained from healthy donors of the San Diego Blood Bank or procured commercially (HemaCare). Peripheral blood mononuclear cells (PBMCs) were isolated by gradient centrifugation (Lymphoprep, StemCell Technologies) and cryopreserved in FBS with 10% DMSO pending further processing.

### Synthesis of barcoding oligonucleotides

Barcoding oligonucleotides all use the following format:

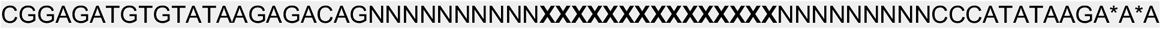

Where **X** represents the barcode sequence, and A* represents a phosphorothioated bond to inhibit nuclease degradation. Barcode sequences are shown in Table S1. Barcode sequence sets were designed with a minimum edit distance of 7 nucleotides from all other barcodes in the set. CellRanger allows a single barcode mismatch, meaning at least 6 sequencing and/or PCR errors would be required for one barcode to be miscounted as another.

### Preparation of next-generation DNA-barcoded antigen baits

Protein antigens encoding a C-terminal Avi-tag [77] were expressed and purified as described previously [49,63,65], and site-specifically biotinylated (BirA500 biotin-protein ligase kit, Avidity LLC) according to the manufacturer’s protocol. Barcoding complexes containing streptavidin (**SAV**), a barcode oligonucleotide and, optionally, a fluorophore were either procured commercially (TotalSeq-C, Biolegend) or created in-house. To create custom antigen barcoding reagents, 5’-biotinylated barcode oligonucleotides (Integrated DNA Technologies) were incubated with streptavidin (ThermoFisher Scientific) or a streptavidin-conjugated fluorophore (AF647 or AF488, ThermoFisher Scientific; BV421, BD Horizon) at a 2.5:1 molar ratio at room temperature for 30 minutes. AgBCs were generated by incubating biotinylated protein antigens with custom or commercially procured barcoding reagents at a molar ratio of either 2:1 or 4:1, depending on the molecular weight of the protein antigen (Table S2). The conjugation reaction was incubated at room temperature and protected from light for a minimum of 30 minutes. Lastly, 200 nM of free unlabeled biotin was added to the AgBCs and left to incubate at room temperature for 15 min to saturate any unoccupied binding sites on streptavidin.

### Cell labeling

Aliquots of cryopreserved PBMCs were thawed by gentle agitation in a 37°C water bath and, when completely thawed, transferred to a 15 mL conical tube containing 10 mL of sterile, pre-warmed post-thaw medium consisting of RPMI 1640 with 50% FBS. Cells were centrifuged at 400g for 5 min, supernatant was discarded, and cell pellets were gently resuspended in 5 mL cold sterile FACS Buffer (1X DPBS with 1% FBS, 1 mM EDTA and 25 mM HEPES). An aliquot of 20 μL of each PBMC sample was set aside for cell counting and verification of viability. The FACS antibody master mix (Table S3) was freshly prepared on ice while protected from light. Aliquots of 10-20 million PBMCs were resuspended on ice with 100 μL master mix along with 200 nM of a “dark” (no fluorophore) HSA (Acro Biosciences) AgBC and 1.25 μg of a unique cell hashing antibody for each sample (TotalSeq-C anti-human or anti-mouse Hashtag, BioLegend). Cells were incubated for 15 min on ice. For experiments involving epitope knockout AgBCs, 200nM of each KO AgBC was added and cells were incubated for an additional 15 min on ice in the dark. Next, 200 nM of the remaining AgBCs were added and cells were incubated for 30 min on ice in the dark. Finally, 1 mL of 1:300 diluted dead cell staining reagent (LIVE/DEAD Fixable Aqua, ThermoFisher Scientific) was added and cells were incubated on ice in the dark for 15 mins. Cells were washed twice with 10 mL of cold FACS buffer, resuspended at the desired concentration using FACS buffer and filtered twice through a 35 μm nylon mesh filter (Falcon) to remove cell aggregates. The second filtration was done immediately prior to sorting.

### Cell sorting and post-sort processing

Prior to sorting, an appropriate number of capture wells in a 96-well PCR plate were prepared by filling with sterile 100% FBS to completely coat the well surface. FBS was then removed from each capture well and 20 μL of fresh FBS was added. Plates were sealed with sterile foil seals (AlumaFoil, Qiagen) and kept at 4°C until the start of the sorting process. When ready to sort, the seal was removed, and the plate was placed onto the pre-chilled plate sample collection platform of a FACSMelody (BD Biosciences). Cells were bulk sorted into the 96-well collection plate using the Purity sort mode and at a flow rate not higher than 1500-2000 events per second to ensure sorting efficiency of 85-95%. Up to 15,000 cells were sorted into a single well. Once sorting was completed, the 96-well capture plate was removed and 80 μL of cold, sterile PBS was added to each capture well to dilute the FBS concentration to 10% v/v or less. The capture plate was again sealed with a sterile foil seal and centrifuged at 2500 rpm for 2 minutes. The seal was removed and 38.6 μL of buffer was added to an adjacent empty well to serve as a reference point for buffer removal. Excess 10% FBS was carefully removed from each sort well until the remaining volume matched the Trypan Blue reference well.

### Single cell library generation

Single cell sequencing libraries were prepared using the Chromium Controller (10x Genomics) by following the provided protocol (*Chromium Single Cell 5’ Reagent Kits User Guide (v2 Chemistry Dual Index) with Feature Barcoding technology for Cell Surface Protein and Immune Receptor Mapping*; CG000330, Rev C) with a few critical modifications:

- **Step 1.2b-c:** Master Mix was added to the pelleted cells (rather than suspended cells being added to the Master Mix). Cells + Master Mix were mixed by very gently pipetting 5-7 times while being careful not to introduce bubbles. Cells + Master Mix were then carefully added to the appropriate well of the Next GEM Chip. This is the ***<u>MOST CRITICAL</u>*** step of the process and should be performed as gently as possible, as any additional cell stress can significantly affect downstream recovery.
- **Step 1.4e:** After completing the Chromium reaction, the resulting emulsions were aspirated *<u>very slowly</u>* (aspiration should take 20 seconds) and *<u>very slowly</u>* transferred into a chilled PCR tube (dispensing should take 20 seconds).
- **Step 3.0e:** If fewer than 1,000 cells were sorted, the total number of amplification cycles was increased to 10.
- **Step 3.3d:** If fewer than 1,000 cells were sorted, the total number of amplification cycles was increased to 10.
- **Step 3.5a:** If the BioAnalyzer trace showed an extra “shoulder” peak adjacent to the expected V(D)J library peak, an additional SPRI cleanup (Step 3.4) was performed.

### Sequencing

The resulting libraries were quantified (Qubit) and pooled at a 5:1 mass ratio between V(D)J and feature barcode libraries, and sequenced with a target depth of at least 10,000 reads per cell (5,000 VDJ reads and 5000 feature barcode reads per cell). Gene expression libraries were sequenced with a target depth of 30,000 reads per cell. Libraries were loaded at 750 pM with 2% PhiX on a NextSeq 2000 using 100-cycle P3 reagent kits or on a NovaSeq 6000 using 100-cycle SP reagent kits with the following run parameters:

- Read 1: 26 cycles
- i7 index: 10 cycles
- i5 index: 10 cycles
- Read 2: 90 cycles

### Data processing

Raw sequencing data was processed using CellRanger (version 6.0.2 or 6.1.2) to generate assembled VDJ contigs and counts matrix files. Briefly, cellranger mkfastq was used to create FASTQ files from sequencing base call (**BCL**) files. We then used cellranger multi to process GEX, VDJ and feature barcode data for each Chromium reaction. Output from CellRanger was read and annotated using scab, as described above. The source code for scab is freely available under the permissive MIT license via GitHub (github.com/briney/scab) and can be installed using the Python Package Index (pypi.org).

Gene expression data were processed using scab and scanpy [53]. Cells were removed if they contained fewer than 200 genes, more than 3,500 genes, or more than 10% mitochondrial gene counts. Genes found in fewer than 0.1% of cells were also removed. Counts were normalized using scanpy’s “cell_ranger” normalization flavor prior to being log plus one transformed. Dimensionality of the gene counts matrix was reduced by performing a principal component analysis, the neighborhood graph was computed, cells were clustered using the Leiden algorithm [78] and the neighborhood graph was embedded using UMAP [79,80]. To minimize the influence of individual immunoglobulin (**Ig**) germline gene segments, which are not directly related to B cell phenotype, Ig germline gene segments are not considered when performing dimensionality reduction, clustering or embedding. Ig genes were removed from consideration if they matched the following regular expression:

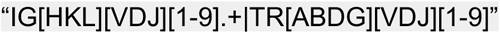

Leiden clusters were assigned to the appropriate B cell developmental subset using genes known to differentiate naive B cells (*IGHD* and *TCL1A*), memory B cells (*CD27, IGHG1-4, IGHA1-2* and *IGHE*), and atypical B cells (*FGR, GDI2, FCRL5*, and *ITGAX*) (Figure S1).

### Calculation of LIBRA-seq recovery efficiency

Recovery efficiency was not reported in the original LIBRA-seq publication, however, the authors did report the number of cells for which paired Ab sequences and antigen specificity data were recovered, as well as the “Targeted Cell Recovery” of each 10x Genomics Chromium reaction. Targeted Cell Recovery is a calculation performed as part of the 10x Genomics Chromium protocol which determines the total number of cells that should be loaded to ensure that the targeted number of emulsion droplets contain both a cell and a barcoded GEM bead (*Chromium NextGEM Single Cell V(D)J Reagent Kits v1*.*1 User Guide with Feature Barcoding technology for Cell Surface Protein*; CG000208, Rev G). We can thus use the Targeted Cell Recovery to back-calculate the total number of input cells and compute overall recovery efficiency for each reaction. In the 10x Genomics protocol, the Targeted Cell Recovery dilution calculation is rounded to the nearest 0.1uL, so the number of input cells may vary slightly based on the starting concentration of input cells. For each Targeted Cell Recovery value, we computed the number of input cells for every starting concentration listed in the 10x Genomics protocol and used the minimum for the following recovery efficiency calculations. For the Ramos B cell lines, 2,321 cells were obtained using a Targeted Cell Recovery of 10,000 cells, indicating a minimum input of 16,740 cells (range: 16,460-16,600) and an overall recovery efficiency of 13.9%. For donor NIAID45, 889 cells were obtained using a Targeted Cell Recovery of 9,000 cells, indicating a minimum input of 14,790 cells (range: 14,790-14940) and an overall efficiency of 6.1%. For donor N90, 1,465 cell were recovered from a Targeted Cell Recovery of 4,000 cells, indicating a minimum input of 6,560 cells (range: 6,560-6,660) and an overall efficiency of 2.3%.

## ACKNOWLEDGEMENTS

This work was funded in part by the Bill and Melinda Gates Foundation, NIAID (UM1 Al1444462 and U19 AI135995), NIGMS (R35 GM133682) and the International AIDS Vaccine Initiative (IAVI) Neutralizing Antibody Consortium (NAC). Jonathan Hurtado, PhD, was supported in part by an NIH T32 training grant to the department of Immunology and Microbiology at Scripps Research (T32 AI007244)

## SUPPLEMENTARY INFORMATION

**Table S1.**
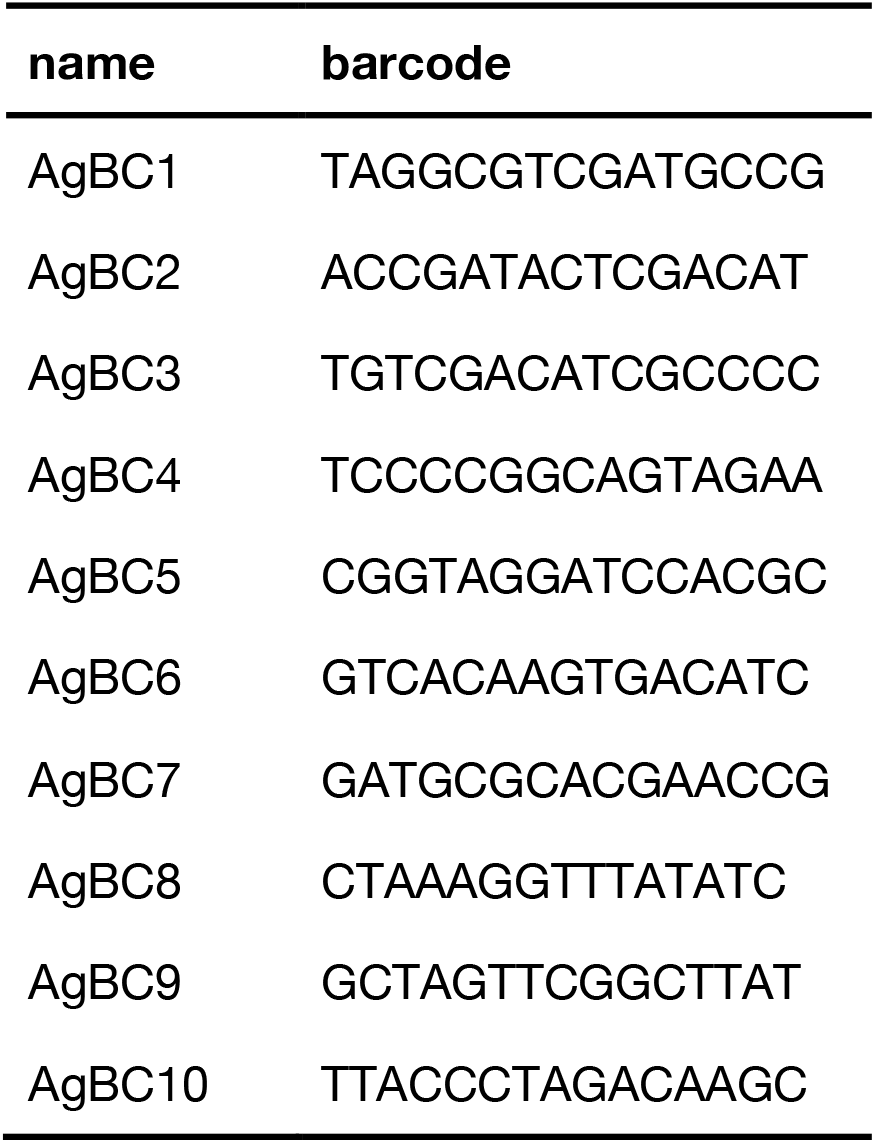
Barcoded oligonucleotides for AgBC construction.

**Table S2.** Preparation of baits based on molecular weight.

|  | **BG505 SOSIP** | **eOD-GT8.1** |
| --- | --- | --- |
| bait concentration | 200 nM | 200 nM |
| molar ratio of bait:SAV | 2:1 | 4:1 |
| bait molecular weight | 225 kD | 22 kD |

**Table S3.** Cell staining master mix (by species).

| **specificity** | **mouse** | **human** |
| --- | --- | --- |
| CD3e | - | APC-Cy7 |
| CD4 | APC-Cy7 | APC-Cy7 |
| CD8a | APC-Cy7 | APC-Cy7 |
| Ly6C | APC-Cy7 | - |
| CD11c | APC-Cy7 | - |
| CD14 | - | APC-Cy7 |
| F4/80 | APC-Cy7 | - |
| CD19 | PE | PerCP-Cy5.5 |
| IgD | PerCP-Cy5.5 | - |
| IgM | BV786 | PE |
| IgG | - | BV786 |

**Figure S1.**
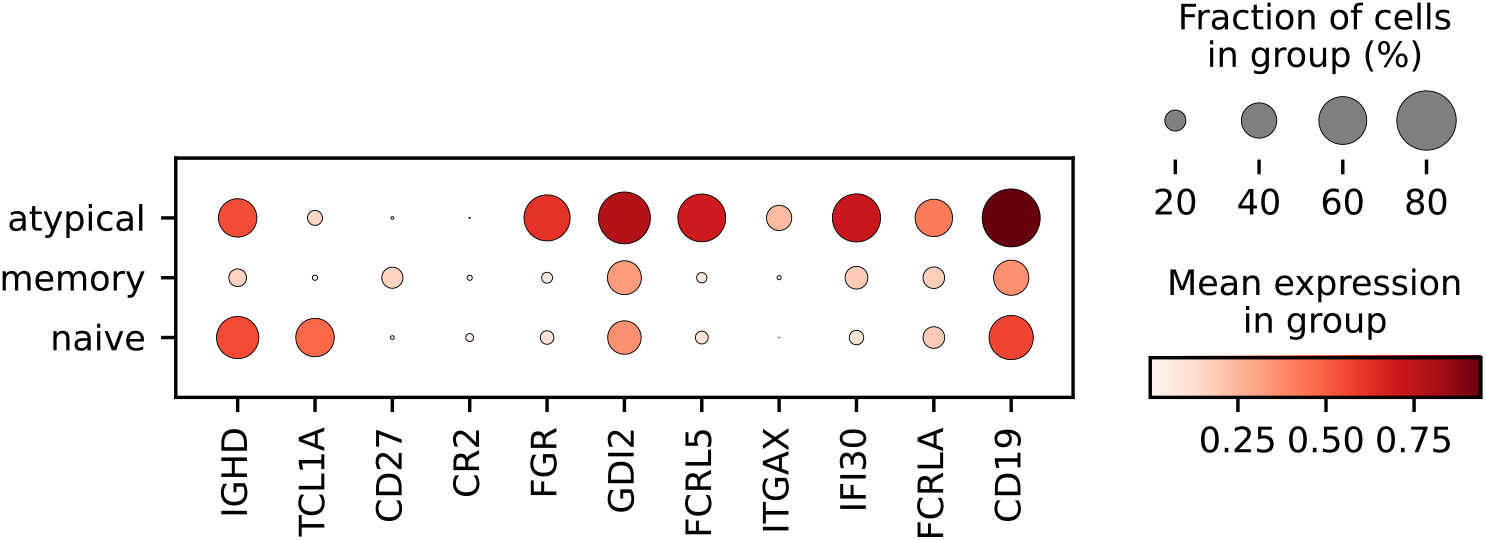
Genes distinguishing naïve, memory and atypical B cell subsets. Plot was created in scanpy using scanpy.pl.dotplot().

